# Evaluation of Phage and Antibiotic Combinations for *Staphylococcus aureus* Biofilm Eradication: Insights from the Wax Worm Model

**DOI:** 10.1101/2023.11.25.568669

**Authors:** L Archana, Prasanth Manohar, Bulent Bozdogan, Ramesh Nachimuthu

## Abstract

**Background:** *Galleria mellonella*, an advantageous model organism, to investigate the therapeutic potential of bacteriophages. Employing the *Galleria mellonella* (waxworm) model, this study involved infecting the larvae with pathogenic *Staphylococcus aureus*, followed by combination treatment with staph phages; vB_Sau_Saa90 and vB_Sau_Saa165 at the concentrations of 10^7^ and 10^9^ PFU/mL and oxacillin, at 100 mg/kg. The treatment modalities were executed in three distinct orders: pre-phage treatment, simultaneous treatment, and post-phage treatment. Each treatment was administered at 90 min intervals. The health state index of the waxworms was monitored throughout the study.

**Results:** Among the three treatment orders, both pre-phage treatment and post-antibiotic treatment exhibited the significant outcomes. The survival percentages ranged from 40% to 50%, 30% to 50%, and 50% to 70% across strong, moderate, and weak biofilm conditions. Notably, the phages vB_Sau_Saa90 and vB_Sau_Saa165 displayed remarkable effectiveness in combination with antibiotics. This successful synergy showcased the potential of combinatorial treatment in effectively addressing biofilm-related infections.

**Conclusions:** This study holds considerable significance as it underscores the suitability of *G. mellonella* as a model organism for exploring bacteriophage-based therapeutic approaches. The study’s findings shed light on the efficacy of combining bacteriophages and antibiotics to target *S. aureus* biofilm cells. The observed results in the waxworm model highlight a promising avenue for managing biofilm infections. However, the need for further validation through larger animal models is emphasized, as this could potentially pave the way for novel treatment strategies with broader clinical implications.

## Introduction

*Staphylococcus aureus* is a Gram-positive bacterium that is known to cause a wide range of infections in humans, including skin infections, pneumonia, endocarditis, and sepsis. *S. aureus* is a common human pathogen that colonizes the skin and nasal passages of up to 30% of the population without causing any symptoms. The emergence of antibiotic-resistant strains of *S. aureus*, particularly MRSA, has become a major public health concern in recent years [1].

Phages are viruses that specifically infect and kill bacteria [2]. Phages have several advantages over antibiotics as a treatment for bacterial infections. They are highly specific, targeting only the bacteria they recognize, and do not harm beneficial bacteria in the body. Phages can also be tailored to target specific bacterial strains or even specific virulence factors [3]. Phage therapy has been successfully used to treat a variety of bacterial infections in humans and animals, including those caused by *S. aureus*. However, there are several challenges associated with the use of phages for therapeutic purposes. These include the potential for the development of phage-resistant bacterial strains, the potential for the transfer of virulence genes between phages and bacteria, and the potential for adverse immune reactions in the host [4].

Animal models are essential tools for studying the safety and efficacy of phage therapy *in vivo*. Traditionally, mammalian models, such as mice and rabbits, have been used to study the pharmacokinetics and toxicity of phages. However, these models are expensive, require specialized facilities, and are subject to ethical considerations. In recent years, non-mammalian animal models, such as *Galleria mellonella* and *Caenorhabditis elegans*, have emerged as attractive alternatives for *in vivo* studies of bacterial infections and phage therapy. *G. mellonella*, also known as the wax moth larva, is a widely used model organism for studying bacterial infections and immune responses. *G. mellonella* larvae have innate immune systems that share many similarities with those of mammals, including the production of antimicrobial peptides and the activation of phagocytic cells [5]. Moreover, *G. mellonella* is an ideal model for high-throughput screening of potential antimicrobial agents, including phages, due to its small size, short lifespan, and ease of handling [6]. Previous studies have demonstrated the efficacy of phage therapy against *S. aureus* infections in *G. mellonella* larvae [7] [8]. Previously, Galleria mellonella has been used to test the phage efficacy against *E. coli, Citrobacter sp*., *Enterobacter sp*., *Klebsiella pneumoniae* [9-11]. *In vivo* studies using animal models can provide valuable insights into the safety and efficacy of phage therapy. In particular, the use of non-mammalian animal models, such as the wax moth larva has gained popularity in recent years due to their low cost, ease of handling, and similarity to mammalian hosts in terms of innate immune responses.

In this research paper, we aim to investigate the *in vivo* efficacy of phage therapy against *S. aureus* biofilm cells using *G. mellonella* as model organisms. Specifically, we explored the ability of phages to improve survival in infected larvae and nematodes.

## Materials and Methods

### Bacteria, bacteriophage, and culture conditions

A clinical *S. aureus* strain, SA-90 and SA-165 and their respective phage vB_Sau_S90 and vB_Sau_S165 used in this study were previously reported [12]. Bacterial isolate SA-90 and SA-165 was originally collected from a diagnostic laboratory in Chennai, India [13]. The isolates were obtained from a pus sample, and the antibiotic resistance analysis by PCR showed that the isolate harbored the *mec*A gene, and was thus classified as methicillin-resistant *S. aureus* (MRSA). Bacterial cultures were maintained in Brain Heart Infusion agar (HiMedia, India) at 37 □for all the analysis.

### Phage isolation and characterization

Phages were isolated by enrichment technique from hospital sewage sample. Characteristic features of the phage, including its life cycle and time-kill analysis, and morphological analysis by Transmission Electron Microscopy has been reported previously [12]. *In vitro* phage-antibiotic synergy was performed as reported earlier [14].

### Study setup and treatment strategies

The efficacy of oxacillin and phage combination *in vivo* was assessed in this study in *G. mellonella* models against biofilm cells. The eradication of bacterial biofilm cells was assessed using the phage and antibiotic combination. The treatment order, otherwise known as the administration of antibiotics and phage at the different points was assessed. Bacterial eradication by the phage and oxacillin combination was assessed using three treatment strategies: pre-phage and post-antibiotic, simultaneous phage and antibiotic, and pre-antibiotic and post-phage treatments.

The treatments were assessed using phages vB_Sau_S90 and vB_Sau_S165. Each treatment strategy was performed independently with phage in combination with oxacillin antibiotic. The concentrations used for the study were 10^9^ PFU/mL of phage in combination with antibiotic. In the pre-phage and post-antibiotic treatment, the bacterial cells were initially treated with the phage for 90 mins, followed by the addition of a concentration of oxacillin treatment for the following 24 hrs. For simultaneous phage and oxacillin treatment, both phages and antibiotics were administered simultaneously and incubated for 24 hrs. For pre-antibiotic and post-phage treatment, the bacterial cells were initially treated with the sub-inhibitory concentrations of oxacillin for 60 mins, followed by phage treatment for the following 24 hrs. The bacterial reduction was measured at OD_600nm_ every 2 hrs for 24 hrs. The experimental setup used for *G. mellonella* is shown in Fig. 1.

**Figure 1:**
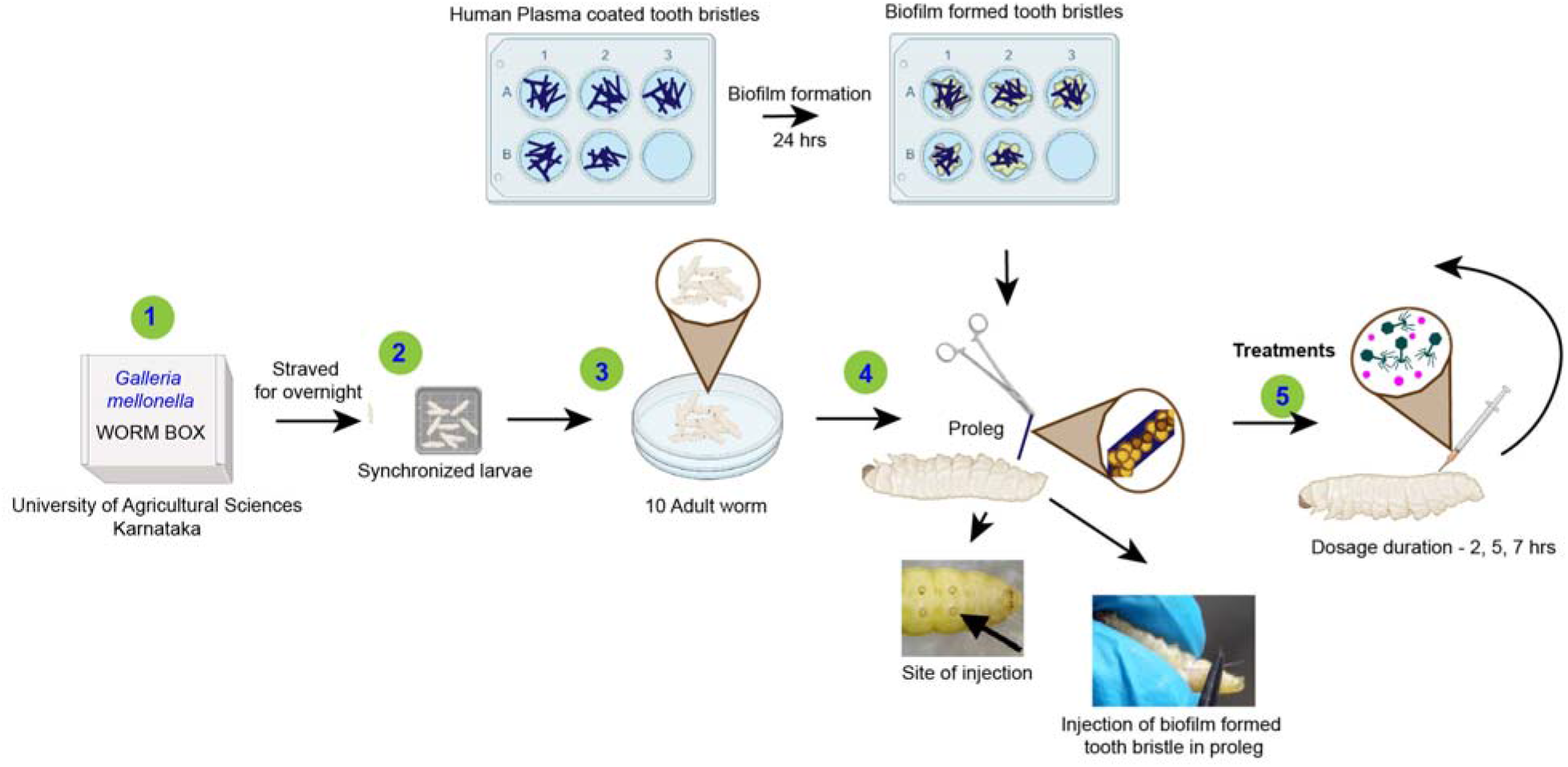
Schematic diagram of experimental setup used for *G. mellonella* biofilm model. 1. *G. mellonella* was harvested. 2. The adult (creamy white color) worms were chosen and starved for 24 hrs. 3. The starved worms were separated into different groups each consisting of 10 worms. 4. The tooth bristle with biofilms are injected into the proleg of experimental worms. Note: The biofilm coated tooth bristles should be prepared in advance. 5. The infected worms were treated with phage-antibiotic combinations. In this study three distinct orders were used i.e., pre-phage treatment, simultaneous treatment, and post-phage treatment at 90 min intervals.

### *In vivo* efficacy in *Galleria mellonella*

*In vivo* biofilm eradication was assessed in *G. mellonella* by introducing biofilm-coated tooth bristles and treating them with phage and oxacillin antibiotic combination. *G. mellonella* at the instar larval stage was collected from the Department of Entomology, University of Agriculture Sciences, Bangalore, India. The received larvae were maintained at the Antibiotic Resistance and Phage Therapy Laboratory, VIT, Vellore. The worms were fed, and supplements were changed every 5 days. Worms were reared and maintained according to the protocol described earlier [15]. Larvae were reared in a dark and maintained at 15–20 °C.

The bacterial isolates SAA.28 (strong biofilm producer), SAA.90 (moderate biofilm producer), and SAA.165 (weak biofilm producer) were used to assess the phage efficiency. For this study, creamy white adult larvae of the fifth instar stage, 20 mm in length and 300–500 mg in weight, were selected. Each study group consisted of ten larvae, which were starved for 20–24 hrs before treatment initiation. Treatments were assessed using oxacillin antibiotic as they showed greater efficiency than other antibiotics assessed in combination with phages *in vitro* [14]. Briefly, for the biofilm formation, the overnight culture was diluted to 1:10 with tryptic soy broth supplemented with 1% glucose and added to 16-well titer plates containing sterilized tooth bristles coated with 10% human plasma and incubated for 24 hrs. The tooth bristle was recovered from the titer plate wells, washed (to remove planktonic cells) and injected aseptically into the left proleg of the worm. The treatment was initiated after 2 hrs with a dosage duration of 2^nd^-, 5^th^-, and 7^th^-hr intervals. For each treatment strategy, phages vB_Sau_S90 and vB_Sau_S165 were combined with oxacillin antibiotic at 100 mg/kg body weight. The control groups were maintained as biofilm-treated, phage-treated, and antibiotic alone treated groups.

The health state of the worms was observed every 24 hrs for 5 days. The efficiency of the treatment was determined based on the larvae’s health using four major health index activities: cocoon formation, melanization, activity, and survival. The health index score was calculated by observational activity, with dead worms marked as 0, and cocoon formation was considered healthier and scored as 1. The efficiency of the treatment strategies and phage combinations was assessed in comparison with that of the control groups. The percentage of survival was calculated for the treated and untreated wax worms. The health state and the progression of infection in the worm is shown in Fig. 2.

**Figure 2:**
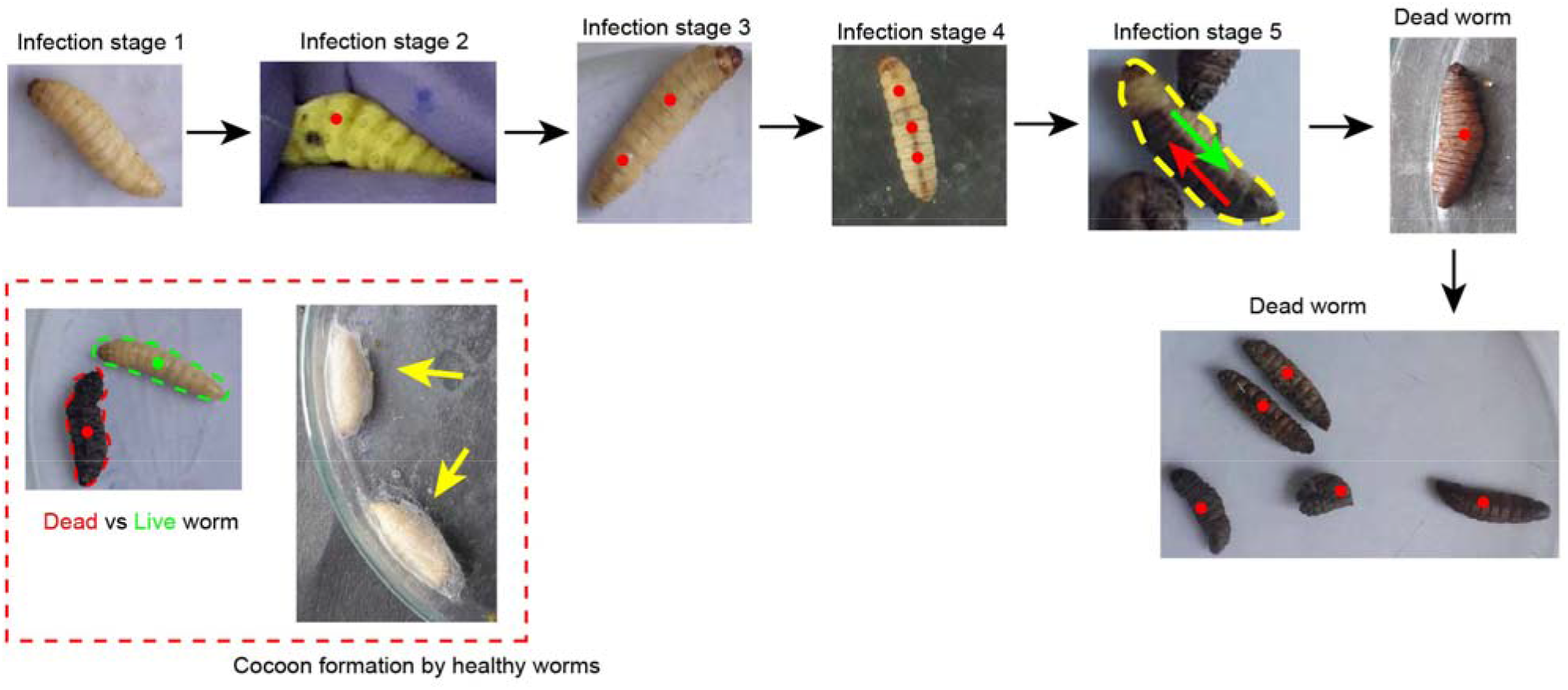
Different stages of infected *G. mellonella* worms; live/ healthy worms-creamy white colored and infected/ dead worms-black spots/ black colored. Stage 1: Infected healthy worms, Stage 2: Infected worms with black spots, Stage 3 & 4: Infected worms with more than one black spots, Stage 5: Infected dead worms turning black. Cocoon-formation denotes the healthy worms.

## Results

### Phage efficacy against biofilm cells in *Galleria mellonella*

The wax worm model was used to study the eradication of the biofilm. Three different levels of biofilm assessed showed that all the biofilm-treated wax worms without any treatment had a 0% survival on 2^nd^ day. The strong biofilm producers (SAA.28) infected wax worms showed that treatment controls, antibiotic alone showed 0% survival on 3rd day, while phage alone treatment showed 10% and 20% survival with phage vB_Sau_Saa90 and vB_Sau_Saa165, respectively. The phage and antibiotic combinatorial treatment showed higher efficiency when compared to phage only treatment method. Among the three treatment strategies, pre-phage and post-antibiotic treatment showed the highest survival percentage of 40% and 50% with phage vB_Sau_Saa90 and vB_Sau_Saa165, respectively. Followed by pre-antibiotic and post-phage treatment and simultaneous phage-antibiotic treatment, showing closer outcomes ranging between 20% to 30% survival for both phages.

For moderate biofilm (SAA.90), oxacillin antibiotic control showed 0% survival on day 3, while phage-only treatments showed 30% and 40% survival with phage vB_Sau_Saa90 and vB_Sau_Saa165, respectively. The phage and antibiotic combinatorial treatment showed higher efficiency when compared to phage-only treatment. Among the phage-antibiotic combination, pre-phage and post-antibiotic treatment showed the highest survival percentage of 80% and 60% with phage vB_Sau_Saa90 and vB_Sau_Saa165, respectively. Followed by pre-antibiotic and post-phage treatment and simultaneous phage-antibiotic treatment, showing closer outcomes ranging between 30% to 50% survival for both phages.

For weak biofilm (SAA.165), oxacillin antibiotic control showed 0% survival on day 3, while phage-only treatments showed 50% and 60% survival with phage vB_Sau_Saa90 and vB_Sau_Saa165, respectively. The phage and antibiotic combinatorial treatment showed higher efficiency when compared to phage-only treatment. Among the phage-antibiotic combination, pre-phage and post-antibiotic treatment showed the highest survival percentage of 80% and 90% with phage vB_Sau_Saa90 and vB_Sau_Saa165, respectively. Followed by pre-antibiotic and post-phage treatment and simultaneous phage antibiotic treatment, showing closer outcomes ranging between 50% to 70% survival for both phages. The survival graph plot for the treatment of biofilm cells in *Galleria mellonella* is shown in Fig. 3.

**Figure 3:**
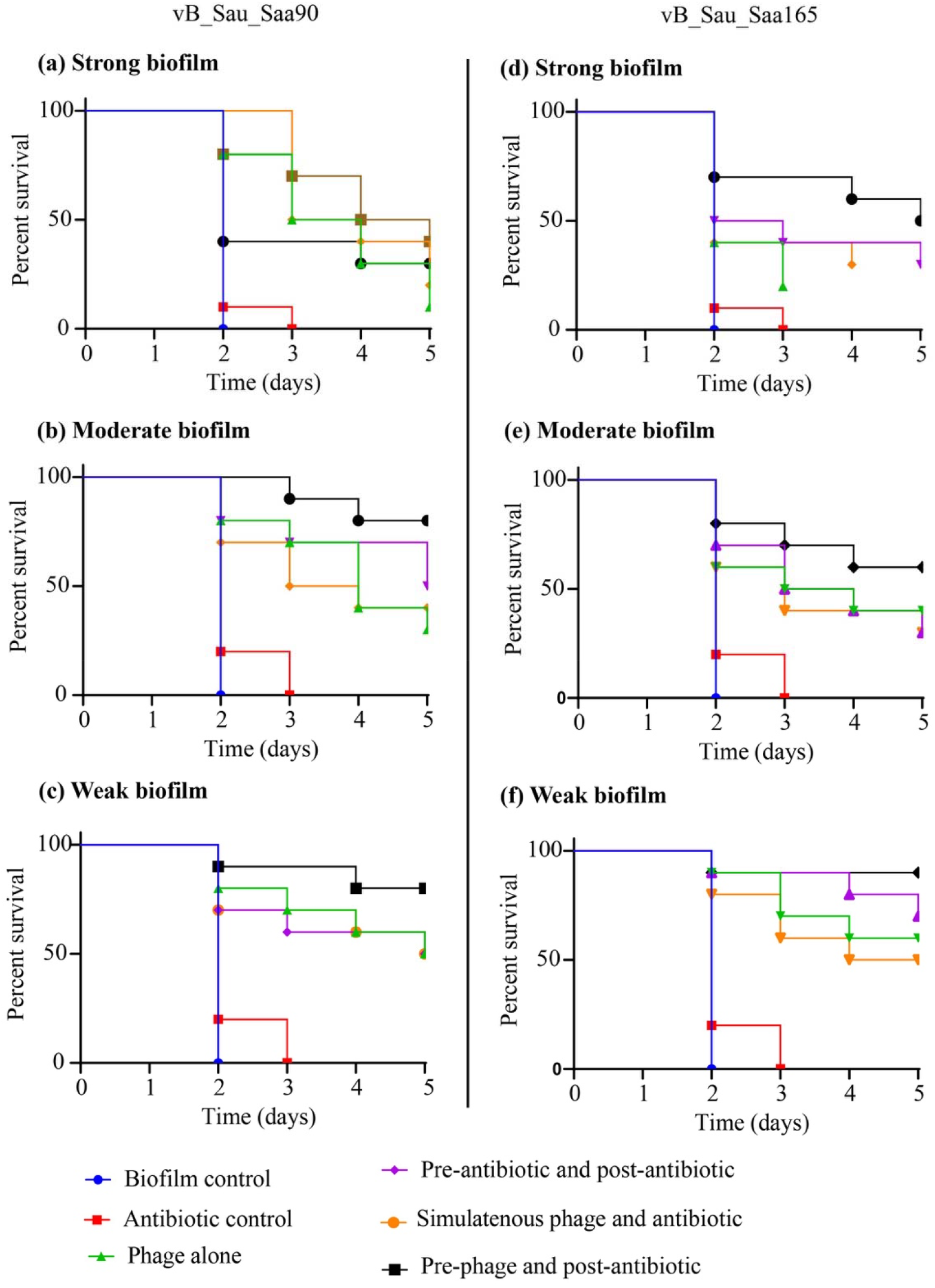
Impact of phage-antibiotic combinations on survivability of *G. mellonella* that was infected with *S. aureus* biofilms and the effectiveness of different strategies (pre-phage and post-antibiotic, simultaneous phage and antibiotic, and pre-antibiotic and post-phage treatments.) of phage-antibiotic combinations. The treatment was initiated after 2 hrs with a dosage duration of 2^nd^-, 5^th^-, and 7^th^-hr intervals. For each treatment strategy, phages vB_Sau_S90 and vB_Sau_S165 were combined with oxacillin antibiotic at 100 mg/kg body weight.

## Discussion

Biofilm-associated infections pose a significant challenge in clinical settings due to their inherent resistance to traditional antibiotic therapies. The use of bacteriophages (phages) as biofilm-targeting agents has gained attention as a promising alternative treatment strategy. This study aimed to evaluate the efficacy of phage and antibiotic combinations in eradicating biofilms using the wax worm model. The results demonstrated the superior effectiveness of the phage-antibiotic combination compared to phage-only or antibiotic-only treatments.

The wax worm model, *Galleria mellonella*, has been increasingly utilized as an *in vivo* model to study microbial infections and evaluate therapeutic interventions. The use of this model provides several advantages, including its cost-effectiveness, ethical considerations, and similarities in immune response to higher organisms, making it a suitable surrogate for studying biofilm-related infections [16].

In this study, the examined biofilms of different strengths and assessed the survival rates of biofilm-treated wax worms under various treatment conditions. The results indicated that without any treatment, the survival rate of wax worms exposed to biofilms was zero on the second day. This emphasizes the robustness and virulence of the biofilm structures.

Phage-only treatments showed varying degrees of efficacy depending on the strength of the biofilm. For strong biofilms (SAA.28), phage-only treatment resulted in 10% and 20% survival rates with phage vB_Sau_Saa90 and vB_Sau_Saa165, respectively, on the third day. For moderate biofilms (SAA.90), the survival rates increased to 30% and 40% with the corresponding phages. Similarly, weak biofilms (SAA.165) demonstrated survival rates of 50% and 60% with the respective phages. These findings are consistent with previous studies that have shown the ability of phages to lyse biofilm-associated bacteria [17, 18]

However, the combination of phages with antibiotics exhibited superior efficacy compared to phage-only treatment. The presence of antibiotics in the treatment regimen likely contributes to the enhanced bacterial killing by disrupting bacterial cell walls, rendering them more susceptible to phage infection. This synergistic effect was evident across all biofilm strengths evaluated in this study. Among the different combinations tested, the pre-phage and post-antibiotic treatment strategy consistently yielded the highest survival rates. This approach involves administering the phage prior to the antibiotic, allowing the phages to initiate bacterial lysis, followed by the antibiotic treatment to further suppress bacterial growth. This sequential administration may facilitate better biofilm penetration and improve bacterial killing efficacy. Similar findings have been reported in previous studies investigating phage-antibiotic combinations against biofilms [19] [20].

These results have significant clinical implications for the treatment of biofilm-associated infections. By combining phages and antibiotics, it may be possible to overcome the limitations of antibiotic resistance and achieve more effective biofilm eradication. Phage therapy offers the advantage of specifically targeting the pathogenic bacteria while minimizing disruption to the commensal microbiota. Furthermore, the use of phages may help overcome the adaptive resistance mechanisms that biofilm-forming bacteria often develop against antibiotics [21].

This study provides valuable insights into the efficacy of phage and antibiotic combinations for biofilm eradication using the wax worm model. The results demonstrate the superiority of the phage-antibiotic combination over phage-only or antibiotic-only treatments. The pre-phage and post-antibiotic treatment strategy showed the highest survival rates across different biofilm strengths. These findings highlight the potential of phage therapy.

The findings from our study using the wax worm model provide valuable insights into the efficacy of different treatment strategies for eradicating biofilms. The results demonstrate that biofilm-treated wax worms without any treatment had a 0% survival rate by the 2nd day, indicating the detrimental impact of biofilms on host survival. This underscores the urgency to develop effective interventions for biofilm-associated infections.

Our results align with previous studies that have investigated the challenges posed by biofilms and the limited effectiveness of antibiotics alone in combating biofilm-related infections. The treatment controls with antibiotics alone showed 0% survival on the 3rd day for both strong (SAA.28) and moderate (SAA.90) biofilm producers. This confirms the need for alternative approaches to overcome the antibiotic resistance mechanisms inherent in biofilms.

Phage therapy, which utilizes bacteriophages to target and lyse specific bacterial strains, has emerged as a potential strategy for biofilm eradication. The survival rates observed with phage-only treatments in our study are consistent with previous literature. For strong biofilm producers, phage-only treatments resulted in 10% and 20% survival with phages vB_Sau_Saa90 and vB_Sau_Saa165, respectively, while for moderate biofilms, the survival rates were 30% and 40% with the same phages.

These outcomes corroborate studies such as Leung and Weitz (2017), which mathematically modelled the synergistic elimination of bacteria by phages and the innate immune system. Their findings suggest that phages can enhance the immune response by reducing the overall bacterial load. This synergistic interaction between phages and the immune system may explain the improved survival rates observed in our phage-only treatment groups [22].

Furthermore, our results highlight the enhanced efficacy of phage-antibiotic combination treatments compared to phage-only treatments. The pre-phage and post-antibiotic treatment strategy demonstrated the highest survival percentages for both strong and moderate biofilms. These findings are consistent with previous studies by Doolittle et al. (1996), which demonstrated the ability of phages to penetrate biofilms and effectively lyse bacterial cells [23]. The sequential application of phages followed by antibiotics in our study likely facilitated biofilm disruption and synergistic bactericidal effects, leading to improved host survival rates.

The observed differences in survival rates among the various treatment strategies for different biofilm strengths (strong, moderate, and weak) emphasize the complexity of biofilm-associated infections and the need for tailored treatment approaches. Our results are in line with the study by Yin et al. (2019), which highlighted the significance of early and accurate biofilm detection methods. Combining such detection methods with phage-antibiotic combinations, as demonstrated in our study, may enable targeted and timely interventions, resulting in improved outcomes for patients [24].

## Conclusion

In conclusion, our study utilizing the wax worm model provides important insights into the efficacy of different treatment strategies for biofilm eradication. The results validate previous literature regarding the limited effectiveness of antibiotics alone and highlight the potential of phage therapy in targeting biofilms. The superior efficacy of phage-antibiotic combination treatments, as observed in our study, underscores the importance of optimizing treatment regimens for biofilm-associated infections. These findings contribute to the growing body of evidence supporting the potential of phage therapy as an adjunctive approach to overcome the challenges posed by biofilms, ultimately improving patient outcomes.

## Acknowledgments

The authors would like to thank Vellore Institute of Technology for providing the necessary facilities to carry out this work. The author, AL gratefully acknowledges the Indian Council of Medical Research (ICMR) for providing financial assistance in the form of Senior Research Fellowship.

## Author Contributions

Conceptualization, A.L., P.M. B.B. and R.N.; Methodology, A.L., P.M., and R.N.; Software, A.L.; Validation, A.L., and R.N.; Formal Analysis, A.L. and P.M.; Investigation, R.N.; Resources, R.N.; Data Curation, A.L.; Writing – Original Draft Preparation, A.L.; Writing – Review & Editing, P.M., B.B. and R.N.; Funding Acquisition, R.N.” All authors have read and agreed to the published version of the manuscript.

## Data Availability Statement

All data are represented as figures, tables. Any further inquiries may be referred to the corresponding author.

## Ethical Statement

Ethical review and approval by the Institute, Vellore Institute of Technology (VIT/IAEC/2023).

